# Virtual reality reveals fine-scale alterations in behaviour following loss of the ADHD-linked gene adgrl3.1 in zebrafish

**DOI:** 10.64898/2026.04.20.719162

**Authors:** Phoebe Reynolds, Emily Read, Caitlin Daly-East, Matthew O. Parker, Robert Hindges

## Abstract

Attention deficit/hyperactivity disorder (ADHD) is a neurodevelopmental disorder characterised by patterns of inattention, hyperactivity, and impulsivity. Understanding the genetic and behavioural mechanisms underlying ADHD requires model systems that enable detailed characterisation of behaviour. Zebrafish have emerged as a prominent model for high-throughput phenotypic screening of candidate risk gene mutations for several disorders, including ADHD. However, conventional behavioural tests for ADHD, such as the forced light/dark assay, quantify basic locomotor activity, limiting the characterisation of behavioural differences beyond overall activity levels. Closed-loop virtual reality (VR) assays offer richer data on animal locomotion by linking visual feedback to an animal’s own movements, allowing the investigation of behavioural responses when sensory input is dependent on locomotion. Here, we have used a closed-loop VR paradigm to assess the behaviour in *adgrl3.1* mutant zebrafish, an established model for ADHD. Our results show that mutants exhibit a higher baseline excitability and a lower threshold for initiating motor events, demonstrating that behavioural differences can be characterised in an interactive environment. More generally, these findings highlight the value of closed-loop VR as a complementary tool for investigating gene disruption and establish a scalable platform for more detailed phenotypic screening of candidate gene mutations associated with neurodevelopmental disorders.

## Introduction

Attention-deficit/hyperactivity disorder (ADHD) is one of the most prevalent neurodevelopmental disorders in children and adolescents, impacting approximately 3-5% of the population worldwide (Castelpietra et al., 2022; Erskine et al., 2013; Geissler & Lesch, 2011; Nigg et al., 2020). It is characterised by persistent symptoms of inattention, hyperactivity, and impulsivity, and can cause profound challenges to cognitive, social, and emotional functioning (Acosta et al., 2004). Beyond its core symptomology, individuals with ADHD also frequently exhibit impairments in executive functioning and decision-making (Mowinckel et al., 2015), as well as increased risk of comorbid psychiatric disorders (Katzman et al., 2017). These include anxiety (Quenneville et al., 2022), bipolar disorder (Nurnberger et al., 2014), depression (Powell et al., 2021), and substance use disorders (Arcos-Burgos et al., 2019).

ADHD is highly heritable, with genetic factors estimated to contribute up to 80% of the variance in risk (Faraone & Larsson, 2019; Yadav et al., 2021). Large-scale genome-wide association studies (GWAS) and gene mutant analyses have identified candidate risk genes that may contribute to susceptibility of ADHD, highlighting its polygenic architecture (Demontis et al., 2023). Among these, genes encoding synaptic adhesion molecules, critical regulators of neural development, synaptic plasticity, and circuit specificity, have gathered increased attention (Burbach & Meijer, 2019; Sudhof, 2018). One gene of particular interest is *ADGRL3* (formerly Latrophilin-3, *LPHN3; (Bruxel et al., 2021)*), which encodes an adhesion G protein-coupled receptor involved in synapse formation, cell adhesion, and plasticity, and is highly expressed in brain regions that govern attention, impulse control, and reward processing (Regan et al., 2021; Sando et al., 2019). Variants of *ADGRL3* have been associated with increased ADHD susceptibility, symptom persistence into adulthood, and variations in treatment responsiveness (Arcos-Burgos et al., 2010; Bruxel et al., 2021; Fallgatter et al., 2013; Kappel et al., 2019; Ribases et al., 2011; Vidal et al., 2022). Interestingly, evidence suggests functional disruptions in ADGRL3 may influence dopaminergic signalling, a pathway central to ADHD (Fontana et al., 2023; Lange et al., 2012; Vidal et al., 2022). Despite these strong associations, the extent to which ADGRL3 variation contributes to behavioural phenotypes, and the mechanistic pathways underlying these effects, remain unclear.

Zebrafish (*Danio rerio*) are a powerful model for investigating the neurogenetic underpinnings of ADHD. They have high fecundity, a well-characterised genome, conserved neurobiological pathways, and ethologically relevant quantifiable behavioural repertoires that make them suitable for studying gene-behaviour interactions (Fontana et al., 2025). Consistent with this, zebrafish mutants lacking *adgrl3.1*, the orthologue of human *ADGRL3*, exhibit core ADHD-like phenotypes, including hyperactivity in larvae, and impulsivity and attention deficits in adults (Fontana et al., 2023; Lange et al., 2012; Regan et al., 2021; Sveinsdottir et al., 2023). However, larval ADHD-related phenotyping has relied largely on the forced light/dark (FLD) locomotion test, here abrupt changes in illumination cause measurable changes in swimming activity (Hillman et al., 2025). These assays are widely used to quantify overall locomotor response and their temporal dynamics and can give insight into anxiety-like states (Burgess & Granato, 2007; Luchtenburg et al., 2019; Peng et al., 2016). However, locomotor changes alone do not necessarily identify the underlying behavioural process, and it is difficult to determine whether increased movement reflects hyperactivity or anxiety-like responses (Blaser et al., 2010; Fontana et al., 2022). This distinction is relevant given the high comorbidity between ADHD and anxiety disorders, estimated at 25–40% in clinical populations (Koyuncu et al., 2022). In addition, conventional FLD assays typically present sensory stimuli independently of the animal’s ongoing movements. Consequently, they provide limited insight into how behavioural responses are regulated when sensory feedback changes as a consequence of the animal’s own actions.

To address this relationship between locomotion and sensory feedback, we used a closed-loop virtual reality assay in freely swimming larval zebrafish (Bahl & Engert, 2020; Reynolds et al., 2026; Slangewal et al., 2026). Here, the visual environment changes in response to the animal’s own movements, creating an interaction based on the animal’s real-time position and behavioural responses. This differs from conventional light/dark assays, in which changes in environment are independent from swimming behaviour. By making visual feedback conditional on movement, the VR assay allows us to examine visuomotor responses under different stimulus demands within an interactive environment. In addition, it allows precise measurements of cognitive traits including attentional, impulsive, and innate decision-making behaviours (Bahl & Engert, 2020; Slangewal et al., 2026). The approach therefore complements conventional locomotor assays as we can investigate ADHD relevant behaviours under movement-contingent visual feedback, rather than relying solely on responses to externally imposed illumination changes.

Here, we applied this paradigm to *adgrl3.1* mutant larvae to test whether the effects of mutation extend beyond altered locomotion to differences in locomotor organisation within an interactive environment. By combining detailed behavioural quantification with an interactive visual environment, we aimed to characterise the behavioural consequences of *adgrl3.1* disruption in greater detail. Our findings provide new insight into the role of *adgrl3* in early neurodevelopment and establish the utility of closed-loop behavioural analysis as a scalable, translational platform for the investigating gene-function relationships relevant to ADHD.

## Material and Methods

### Zebrafish

Zebrafish husbandry and experiments were approved by the Animal Welfare and Ethics Review Body at KCL and conducted under the Home Office project license PP7266180 to R.H. Embryos were collected from large group breedings (Facility Mass Embryo Production system) of adults between 6 -18 months in age. Following collection and injection, embryos were transferred and placed into Danieau solution (58 mM NaCl, 0.7 mM KCl, 0.4 mM MgSO₄, 0.6 mM Ca(NO₃)₂, 5 mM HEPES, pH 7.6) within a 90mm Petri dish, at a density of no more than 50 larvae per Petri dish. Zebrafish were maintained in an incubator at a temperature of 28.5°C and on a 14-hour ON/10-hour OFF light cycle. For experiments on larvae over 5 days, larvae were fed live rotifers as in line with the KCL Fish Facility feeding protocol. All experiments were performed on the AB wildtype line (ZIRC, 5^th^ Generation) at 7 days post-fertilization (dpf), at which age sex cannot be determined. All batches of embryos were collected before 11am, with 7dpf established from this point after the 7^th^ day after injections.

### Generation of *adgrl3.1* CRISPant mutant larvae

CRISPant F0 mutants were generated following the CRISPR/Cas9 ribonucleoprotein (RNP) injection method as described by (Kroll et al., 2021), with the concentration of guide RNA (gRNA) and Cas9 reduced to one-quarter of the recommended concentrations of the original protocol. Three CRISPR RNA (crRNA) oligonucleotides targeting *adgrl3.1* were designed using the Integrated DNA Technologies (IDT) CRISPR design tool.

For RNP assembly, each crRNA (50 μM) was individually annealed to trans-activating CRISPR RNA (tracrRNA; 50 µM) by heating to 95 °C for 5 minutes, followed by immediate cooling on ice. The resulting gRNAs (15.25 µM) were then incubated with Cas9 protein (15.25 µM) at 37 °C for 10 minutes. Each of the three RNP complex (7.125 µM) was subsequently combined in equal amounts, producing a final working concentration of 2.5 µM for each RNP. Control RNPs were prepared in the same way using the Alt-R® CRISPR-Cas9 Negative Control #1 crRNA (Integrated DNA Technologies) following the same procedure.

Microinjections were carried out by delivering 1nl of either the experimental or control RNP mixture into each AB wildtype embryo when at the 1-cell stage. Injections were always done within the first 20 minutes of the embryo being laid, with developmental stage confirmed by visual inspection of the 1 cell stage as injection occurred. Injected embryos were subsequently maintained under standard husbandry conditions. The resulting F0 larvae are referred to as *adgrl3.1* CRISPant mutant larvae throughout this manuscript. This refers to larvae generated using CRISPR/Cas9 targeting and assessed by genomic PCR, as described below; however, complete loss of ADGRL3.1 protein expression was not independently established.

### Genotyping of *adgrl3.1* F0 mutant zebrafish

To assess genotyping outcomes, six embryos from each injection batch for experimentation were chosen at random for genotyping at 2 dpf. All guide RNA target sites were first validated by sequencing to confirm the intended genomic target within the exons. Genotyping was performed by PCR amplification of exons 6 and 8 of *adgrl3.1*, and large indels were detected by gel electrophoresis (Figure S1A). To assess possible excision of the intervening genomic region, the exon 6 forward primer was paired with the exon 8 reverse primer, amplifying exons 6–8 plus flanking intronic sequences (Table S1).

Genomic DNA was extract from whole embryos using the HOTSHOT method (Meeker et al., 2007). Larvae were lysed in 30 µl 50 mM NaOH at 95 °C for 20 min, cooled on ice for 5 min, and pH neutralised with 1:10 volume of 1 M Tris-HCl (pH 8). Samples were vortexed for 10 s and then centrifuged at 13,000 rpm for 1 min. PCR was carried out in 15 µl reactions using DreamTaq PCR Master Mix (Thermo Fisher Scientific), containing 7.5 µl 2× mix, 5.4 µl nuclease-free H₂O, 0.3 µl of each 10 µM primer, and 1.5 µl DNA template (supernatant form previous step). Cycling conditions were: 98 °C for 30 s; 35 cycles of 98 °C for 10 s, 60 °C for 30 s, 72 °C for 30 s; followed by final 72 °C for 2 min and a 4 °C hold. PCR products were resolved on 1% agarose gels in 1× TAE buffer containing SYBR Safe DNA Gel Stain (1:10,000, Thermo Fisher Scientific). Gels were loaded with 5 µl DNA sample per well and run along 100 bp ladders (New England Biolabs) at 120 V for 30 min. DNA was visualised under UV illumination with a NuGenius Gel Documentation System.

Embryos were classified as mutants if at least one mutation or excision was detected in any of the targeted regions. All six CRISPant embryos sampled from each injection batch showed evidence of mutations (Figure S1B), aligning with (Kroll et al., 2021) reporting that this method produces biallelic knockouts in over 90% of injected embryos. These results support successful targeting of *adgrl3.1* across the sampled batches, but do not establish the exact sequence or extent of mutation in every larva.

### Behaviour experiments in freely swimming larval zebrafish

The behavioural paradigm was performed using a custom closed-loop system, as previously described (Reynolds et al., 2026). At 7 dpf, larvae were individually placed into custom-made circular acrylic dishes, measuring 120mm in diameter, 5mm in height, with a black rim and transparent base covered with diffusion paper. Dishes were filled with 50 ml of filtered fish water and left to acclimate for approximately 30 minutes in the system before experimentation. One computer was connected to 8 cameras and 4 projectors covering 8 dishes, therefore allowing tracking of up to 8 larvae simultaneously at a rate of 90 Hz. Combinations of stimuli were presented randomly. Swim bouts were considered turns rather than straight bouts if they had an absolute orientation change larger than 5 °. Any trial where the animal does not move in the middle of the dish or could not be accurately tracked was removed automatically by the processing script.

### Freely swimming paradigm

A custom visual stimulus system was developed using Panda3D and OpenGL Shader Language (GLSL) to render real-time dynamic grating patterns within the experimental arenas. Each larva was presented with a drifting sine wave grating at a constant speed of 0.9 cm/s, with a randomly assigned phase offset. Gratings moved either to the left or right relative to the larva’s orientation. A total of 28 unique stimuli were generated by crossing stripe widths (wavelengths) of 0.005, 0.01, 0.02, 0.04, 0.08, 0.16, and 0.32 arbitrary units (AU), corresponding to stripe widths ranging from approximately 0.64 mm to 41.14 mm (measured as one full cycle of black and white). Each of these was combined with one of two contrast levels (0.25 or 0.75) and one of two motion directions (leftward or rightward). Each stimulus was presented for 30 seconds, beginning with a 5-second rest period during which a black screen was displayed, followed by 25 seconds of drifting motion. Each fish experienced at least four trials of each stimulus, with the total experimental duration lasting one hour. Throughout the experiment, water temperature was maintained at approximately 26 °C.

### Statistics and reproducibility

Embryos were randomly collected from large group breedings. Injections of both the mutagenic and control RNP complexes were performed across different collections (between 2 and 6) of the same day and then pooled within each genotype. For behavioural testing, larvae were randomly selected from these injected pools and relabelled to conceal genotype identity from experimenters. This process was repeated across multiple days. Genotype information was only disclosed after completion of the experiments, during data analysis.

Injections of both the mutagenic and control RNP complexes were performed across different collections (between 2 and 6) of the same day and then pooled within each group. For behavioural testing, larvae were randomly selected from these injected pools and relabelled to conceal genotype identity from experimenters. This process was repeated across multiple days. Genotype information was only disclosed after completion of the experiments, during data analysis.

Normality was determined through either the Shapiro-Wilk test or manual inspection of Q-Q plots to first chose the appropriate statistical test. Parametric data were analysed with mixed-effects model to account for any missing completely at random (MCAR) values. Sphericity was not assumed, and the Geisser-Greenhouse correction was applied to the stripe width and interaction terms, with post hoc comparisons performed using Šídák multiple comparisons test where appropriate. Comparisons with a single value per larva were analysed with a two-tailed unpaired t-test when both groups were normally distributed, or with a Mann-Whitney U test if not normally distrubted. These analyses were performed in Prism 11 (GraphPad). For right-skewed data (bout duration, interbout interval, and bout length) a gamma (γ) Generalised Linear Mixed Model (GLMM) with a log link function was used. These were fitted using the lme4 package in R (Version 4.6.2). Models included genotype, stripe width and their interaction as fixed effects, and a random intercept for fish ID and batch to account for repeated measurements of each larva across stripe widths and variation. Post hoc comparisons between mutants and controls at each stripe width were performed on estimated marginal means (emmeans package), using Wald z-tests on the log scale with Holm correction for multiple comparisons across the seven stripe widths.

Statistical significance was set at p < 0.05. Sample sizes were chosen to provide a high statistical power of > 0.8 and based on previous studies (Reynolds et al., 2026). For all experiments, a minimum of n = 62 was run.

## Results

To investigate the role of *adgrl3.1* in ADHD-related behaviours using our VR approach, F0 CRISPants were generated at the one-cell stage (see Methods) and tested at 7 days post-fertilization (dpf) in a closed-loop visual stimulation system (Figure 1A). At this developmental stage, the visual system is functionally mature, with retinotectal projections, lamination, and stable synaptic density established (Niell et al., 2004; Robles et al., 2014). By 7 dpf, larvae also exhibit consistent swimming behaviour (Budick & O’Malley, 2000; Fitzgerald et al., 2019), making them well-suited for visual-motor behavioural assays as high-throughput model.

**Figure 1.**
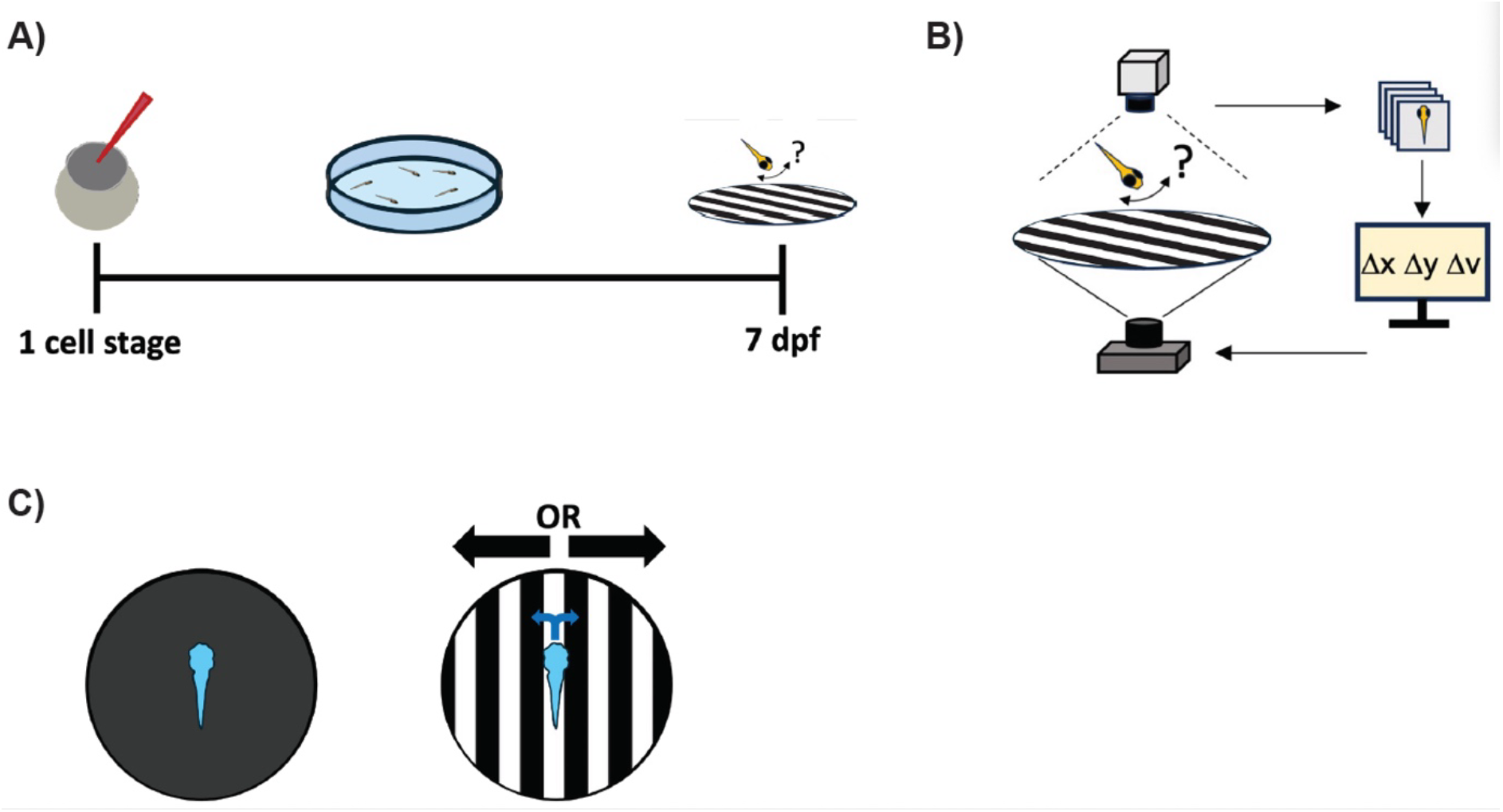
Experimental set up of the virtual reality closed-loop assay. **A)** F0 CRISPant-generated *adgrl3.1* mutants are created at the one cell stage and raised in a petri dish before being assayed at 7 dpf within the closed-loop virtual reality assay. **B)** Diagram of the closed-loop visual stimulation system. Individual larvae are tracked in real time within the arena (12cm diameter), with visual gratings updated and projected such that stimuli remained aligned to the animal’s body axis. **C)** Optomotor response (OMR) stimulus paradigm. Each trial consisted of a 5 second baseline (black screen) followed by 25 seconds of drifting gratings moving either leftward or rightward. Gratings were presented at two contrast levels (0.25 and 0.75 AU) and across multiple spatial frequencies (stripe widths), allowing assessment of contrast- and frequency-dependent visuomotor responses.

Individual larvae were placed into the closed-loop system (Figure 1B), where their position and orientation were tracked in real time. Visual gratings were projected around each larva, with visual stimulus orientation updated to remain fixed on the animal’s body axis, regardless of swimming trajectory. This approach allowed us to assess the visuomotor responses of the larvae under controlled visual stimulation, extending beyond classical locomotor assays conducted in static light/dark environments (Hillman et al., 2024). The stimulus protocol consisted of a 5 second baseline with a black screen, followed by 25 seconds of gratings parallel to the zebrafish larvae moving either to the right or left (Figure 1C). Gratings were presented at two contrasts (0.25 and 0.75 AU) and at multiple stripe widths, as zebrafish responses follow a Gaussian-like dependence on spatial frequency, with a peak at the intermediate widths provided (Reynolds et al., 2026). This allowed us to assess whether loss of *adgrl3.1* alters not only overall locomotor drive, but also specific visuomotor behavioural responses under different stimulus demands.

### Increased locomotion in *adgrl3.1* mutants is independent from increase swim rate and bout duration

To isolate specific parameters during visual motion stimulation, we analysed larval behaviour during the final 25 second of each 30 second stimulus period, during which moving gratings to either the left or right of the larvae’s position were presented. This allowed us to focus on the relationship between locomotion and sensory feedback, rather than possible general anxiety phenotype from drastic changes in luminance. Across all measurements, larvae exhibited the expected Gaussian tuning of swimming behaviours relative the stripe width, with peak responses occurring at approximately 10.3 mm wide stripes, consistent with previous reports (Reynolds et al., 2026) .

Quantification of bout speed showed that mutant larvae swam with significantly higher bout speeds than control-injected larvae (“controls”) under low-contrast stripe conditions (0.25 AU; Figure 2A; Mixed-effects model, Genotype: F(1, 131) = 5.07, P = 0.0259). Post-hoc testing indicated significantly higher speeds in mutants across multiple stripe widths (0.6 mm: P = 0.0225; 5.1 mm: P = 0.0069; 10.3 mm: P = 0.0040; 41.1 mm: P = 0.0317). Importantly, the increase in bout speed in *adgrl3.1* mutant larvae is more pronounced under low conditions than high contrast (0.75 AU) conditions, where overall bout speed did not differ significantly between genotypes (Figure S2A; Mixed-effects model, Genotype: F(1, 130) = 1.93, P = 0.1675), although post-hoc testing indicated significantly higher speeds in mutants at the 5.1 mm stripe width (P = 0.0032).

**Figure 2.**
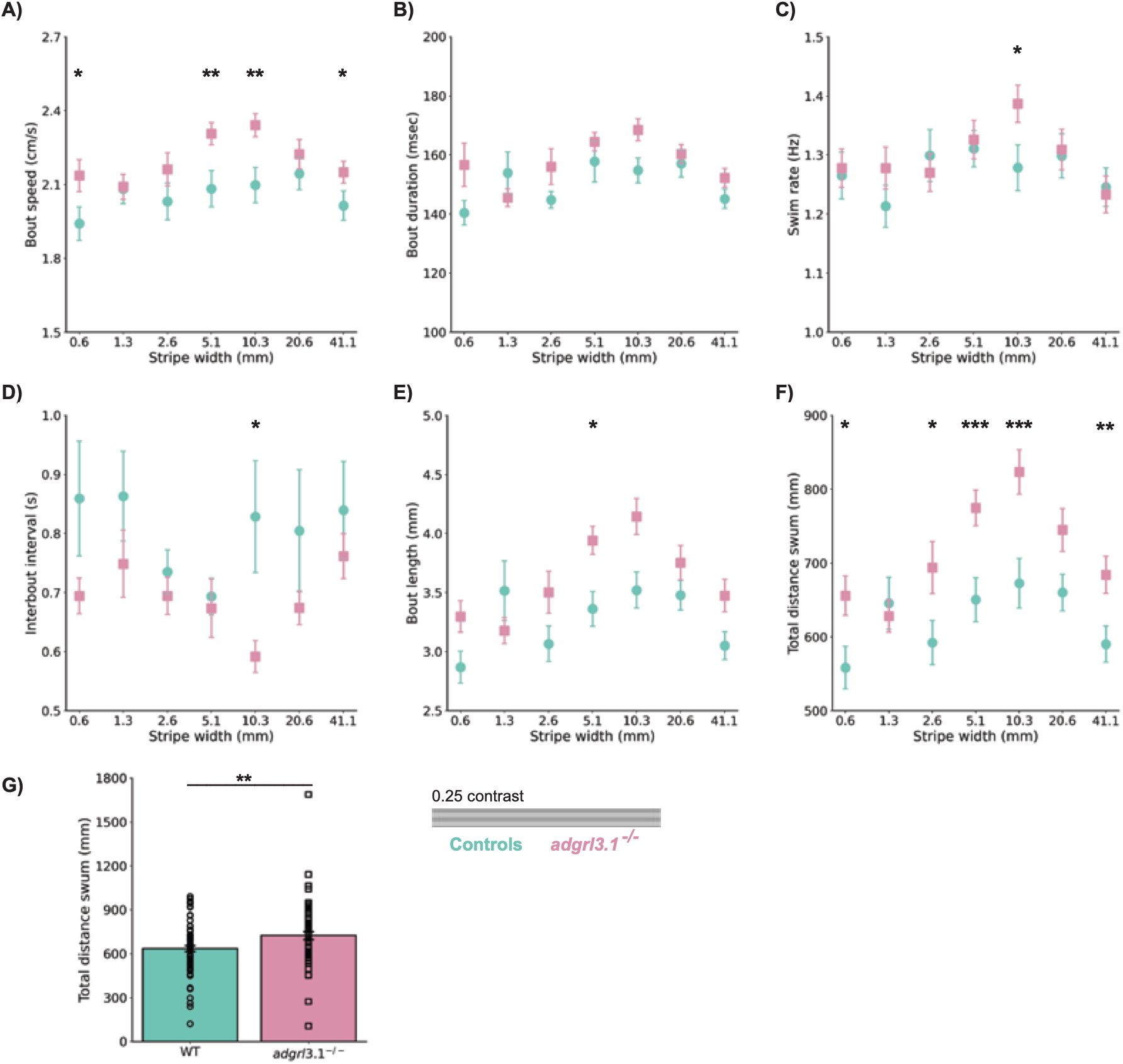
Increased locomotor output in *adgrl3.1* mutants viewing a lower contrast stimulus is driven by altered bout structure. **A-F)** Various swim behaviours measured across stripe widths at a low contrast value of 0.25 AU for control larvae (blue), and mutant *adgrl3.1* mutants (purple). **A)** Average bout speed (Mixed-effects model, Genotype: *P = 0.0259; Stripe width: ***P < 0.001; Stripe width at 0.6 mm: *P = 0.0225; Stripe width at 5.1 mm: **P = 0.0069; Stripe width at 10.3 mm: **P = 0.0040; Stripe width at 41.1 mm: *P = 0.0317). **B)** Average bout duration (γ GLMM, Genotype: P = 0.215; Stripe width: ***P < 0.001. **C)** Swim rate measured in bouts per second (Mixed-effects model, Genotype: P = 0.3613; Stripe width: **P = 0.0079; Stripe width at 10.3 mm: *P = 0.0327). **D)** Interbout interval (IBI) (γ GLMM, Genotype: *P = 0.037; Stripe width: ***P < 0.001; Stripe width at 10.3 mm: *P = 0.017. **E)** Bout length (distance travelled per bout) γ GLMM, Genotype: *P = 0.038; Stripe width: ***P < 0.001). **F)** Total distance travelled (Mixed-effects model, Genotype: ***P < 0.001; Stripe width: ***P < 0.001; Stripe width at 0.6 mm: *P = 0.0151; Stripe width at 2.6 mm: *P = 0.0194; Stripe width at 5.1 mm: ***P < 0.001; Stripe width at 10.3 mm: ***P < 0.001; Stripe width at 41.1 mm: **P = 0.0033). **G)** Total distance travelled at a 0.25 AU contrast condition for control larvae (blue), and mutant *adgrl3.1* mutants (purple) (Mann-Whitney U Test, **P = 0.0093). **A–G)** n = 62 for *adgrl3.1* mutant larvae, n = 70 for control larvae. All error bars are mean ± s.e.m. over fish. Asterisks indicate significance *P < 0.05, **P < 0.01, ***P < 0.001.

We next examined the duration of individual swim bouts (Figure 2B, S2B). Both control and mutant *adgrl3.1* larvae exhibited the expected Gaussian modulation of bout duration across stripe widths, increasing from ∼140 ms at 0.6 mm to ∼170 ms at 10.3 mm. However, no significant main effect of genotype on bout duration was observed under either low-contrast (γ GLMM, Genotype: χ²(1) = 1.54, P = 0.215) or high-contrast conditions (γ GLMM, Genotype: χ²(1) = 1.59, P = 0.207). Similarly, overall swim rate, measured as bouts per second, was not significantly affected by *adgrl3.1* loss under low contrast (Figure 2C; Mixed-effects model, Genotype: F(1, 131) = 0.84, P = 0.3613) or high contrast (Figure S2C; Mixed-effects model, Genotype: F(1, 130) = 0.55, P = 0.4592), with a peak in swim rate at 0.25 contrast (control: 1.2785 Hz ± 0.03899; mutant: 1.3867 Hz ± 0.03147), and at 0.75 contrast (control: 1.4091 Hz ± 0.03439; mutant: 1.4164 Hz ± 0.03712), similar to previous results for larvae in this assay (Reynolds et al., 2026). However, under low contrast, a significant genotype difference was detected specifically at the 10.3 mm stripe width (P = 0.0327).

### *adgrl3.1* mutants exhibit shorter interbout intervals and increased bout length

Next, we concentrated on the assessment of the interbout interval (IBI) and bout distances to determine how *adgrl3.1* loss alters motor initiation and displacement. IBI has been associated with transitions in internal state (Johnson et al., 2020), and therefore, may provide insight into temporal organisation of swimming. Under low contrast, *adgrl3.1* mutants showed a significant reduction in IBI compared to controls, forming an inverse Gaussian curve (Figure 2D; γ GLMM, Genotype: χ²(1) = 4.33, P = 0.037), driven largely by shorter intervals at the 10.3 mm stripe width (P = 0.017) and overall reduced variability in IBI. Under high contrast, this main genotype effect showed a trend toward reduction but did not reach statistical significance (Figure S2D; γ GLMM, Genotype: χ²(1) = 3.03, P = 0.082), although mutants maintained a significantly shorter IBI at the 2.6 mm stripe width (P = 0.013).

Bout length (distance travelled per bout), also revealed that *adgrl3.1* mutants swam significantly longer distances per bout than control larvae under low-contrast stimulation (Figure 2E; γ GLMM, Genotype: χ²(1) = 4.31, P = 0.038). At high contrast, both groups followed a similar pattern, but group overall bout length did not significantly differ between genotypes (Figure S2E; γ GLMM, Genotype: χ²(1) = 1.89, P = 0.169).

Consistent with the increase in bout speed, bout length, and reduced IBIs, the total distance travelled during the stimulus phase was significantly higher in mutants in these low contrast conditions (Figure 2F; Mixed-effects model, Genotype: F(1, 131) = 12.62, P < 0.001), with pairwise elevation across almost all stripe widths (0.6 mm: P = 0.0151; 2.6 mm: P = 0.0194; 5.2 mm: P < 0.001; 10.3 mm: P < 0.001; 41.1 mm: P = 0.0033), reflecting prior studies that measured baseline locomotor activity in *adgrl3.1* knockouts (Fontana et al., 2023; Regan et al., 2021; Sveinsdottir et al., 2023). Mutants consistently swam farther than control larvae, with peaks at 10.3 mm stripe widths both in the lower and higher contrasted stimuli (Controls = 672.9481 mm ± 33.6654; Mutants = 823.5271 mm ± 30.0897). Overall low-contrast total distance collapsed across conditions similarly demonstrated a significant elevation in mutants (Figure 2G;

Mann-Whitney U test, U = 1627, P = 0.009). Under high contrast, total distance also showed a significant overall genotype effect (Figure S2F; Mixed-effects model, Genotype: F(1, 130) = 5.97, P = 0.0159), with specific increases at 2.6 mm (*P* = 0.0044), 5.1 mm (*P* = 0.0039), and 41.1 mm (*P* = 0.0441), as well as when collapse across stripe widths (Figure S2G; Mann-Whitney U test, U = 1734, P = 0.0471). These findings can validate the sensitivity of our closed-loop paradigm to ADHD-related phenotypes, while providing further granular analysis of ADHD-like behaviours.

### *adgrl3.1* mutants display reduced behavioural variability and altered baseline motor structure

To evaluate motor stability and flexibility across genotypes, we quantified the coefficient of variation (CV) of key locomotor parameters, namely IBI and average speed (Figure 3A-B). The CV of the IBI provides a measure of the variability in the timing between consecutive swim bouts, therefore reflecting how flexibly larvae initiate movement across repeated stimuli. Under low contrast conditions, *adgrl3.1* mutants displayed a trend near the significance threshold for average speed CV (Figure 3A; Mixed-effects model, Genotype: F(1, 131) = 3.91, P = 0.0501), with a significant reduction at 10.3 mm (P = 0.0394), while there was reduced IBI variability (Figure 3B; Mixed-effects model, Genotype: F(1, 131) = 5.98, P = 0.0158), with a post-hoc difference also at 10.3 mm (P = 0.0177). Under high contrast, mutants exhibited significantly reduced speed CV across conditions (Figure S3A; Mixed-effects model, Genotype: F(1, 130) = 7.61, P = 0.0067), with post-hoc reductions across multiple stripe widths (0.6 mm, 1.3 mm,2.6 mm, 10.3 mm, 20.6 mm; all *P* < 0.05). High-contrast IBI CV also showed a similar trend (Figure S3B; Mixed-effects model, Genotype: F(1, 130) = 3.80, P = 0.0533; 2.6 mm: P = 0.0161). While the magnitude of movements (i.e. how fast they swim) remains stable across stimuli, this reduction in CV and narrower distributions is reflective of mutants producing swim bouts with greater regularity, resulting in a more stereotyped behavioural profile.

**Figure 3.**
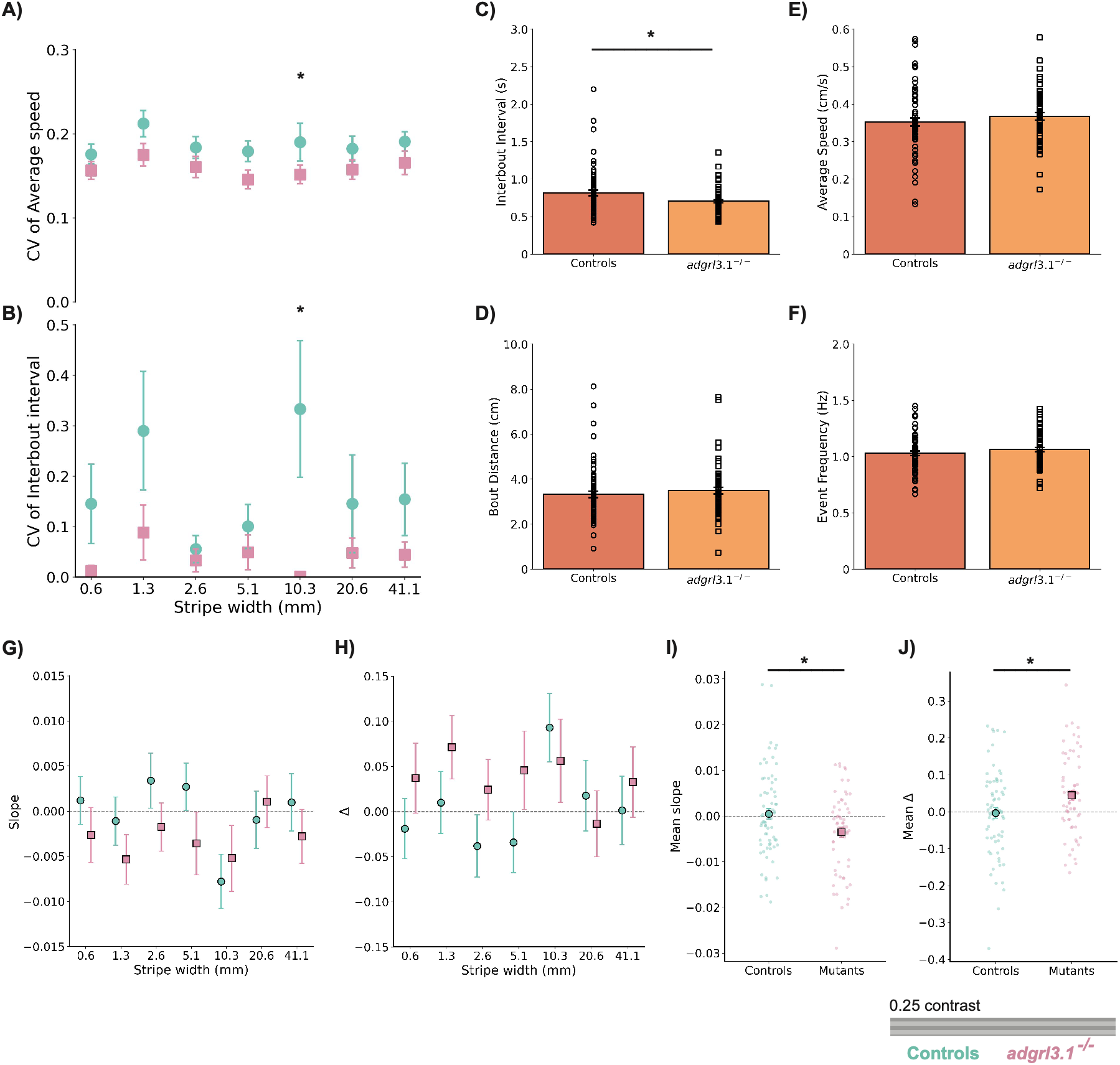
Reduced behavioural variability and altered baseline interbout interval are present in *adgrl3.1* mutants. **A)** Coefficient of variation (CV) for average speed across stripe widths to indicate flexibility in motor output (Mixed-effects model, Stripe width: *P = 0.0299; Genotype: P = 0.0501; Stripe width 10.3 mm: *P = 0.0394). **B)** Coefficient of variation (CV) for interbout interval (IBI) across stripe widths to indicate flexibility in motor output (Mixed-effects model, Stripe width: P = 0.188; Genotype: *P = 0.0158; Stripe width 10.3 mm: *P = 0.0177). **C)** Interbout interval (s) behaviour during the pre-stimulus (black screen) period of each trial (Mann-Whitney U Test, *P = 0.0407). **D)** Bout distance (cm) during the pre-stimulus (black screen) period of each trial (Mann-Whitney U Test, P = 0.2186). **E)** Average speed (cm/s) during the pre-stimulus (black screen) period of each trial (Mann-Whitney U Test, P = 0.3215). **F)** Event frequency, measured as the number of bouts per second during the pre-stimulus (black screen) period of each trial (Mann-Whitney U Test, P = 0.2755). **G)** Within-trial slope of successful optomotor responses over stripe widths. Negative slopes indicate declining performance (vigilance decrement). (Mixed-effects model, Genotype: P = 0.0955; Stripe width: P = 0.2446). **H)** Within-trial delta (Δ) in response success between early and late trial phases across stripe widths, where larger positive deltas indicate decline in performance over trials. (Mixed-effects model, Genotype: P = 0.111; Stripe width: P = 0.374). In G and H, the fish random effect was estimated as zero and removed from the model. **I)** Within-trial slope averaged across stripe widths for each larva (unpaired t-test, t(131) = 2.512, *P = 0.0132). **J)** Within-trial Δ averaged across stripe widths for each larva (unpaired t-test, t(131) = 2.417, *P = 0.0170). **A-J)** n = 62 for *adgrl3.1* mutant larvae, n = 70 for control larvae. All error bars are mean ± s.e.m. over fish. Asterisks indicate significance *P < 0.05.

Before the onset of each visual stimulus, all trials begin with a 5-second black screen during which larvae experienced no visual motion cues. This short pre-stimulus period provided a measure of spontaneous locomotor behaviour in the absence of visual motion cues. Analysing this phase allows assessment of baseline motor excitability, independent of visual feedback, providing a further insight into behavioural phenotypes. Mutants exhibited a significant reduction in baseline IBI compared to controls (Figure 3C; Mann-Whitney U test, U = 1751, P = 0.0407). However, baseline bout distance (Figure 3D; Mann-Whitney U test, U = 1931, P = 0.2186)), average speed (Figure 3E; Mann-Whitney U test, U = 1984, P = 0.3215), and number of bouts (Figure 3F; Mann-Whitney U test, U = 1962, P = 0.2755) did not show significant genotype differences. This selective reduction in baseline IBI indicates an alteration in the temporal regulation of movement initiation prior to sensory stimulation, whereas the overall kinematic scale and movement intensity remain preserved.

### *adgrl3.1* mutants show a within-trial decline in tracking performance under low contrast

Finally, we quantified how visual tracking performance changed over the course of each trial through measuring orienting behaviour to the visual stimuli. Larval orientation was segmented into discrete swim bouts, and movements aligning with the direction of visual motion, with an orientation change > 5°, were classified as successful tracking responses. For each larva and stimulus, the proportion of successful responses was then analysed across sequential time bins within each trial. The within-trial slope represents the linear trend in successful responses over time, with a negative slope indicating declining performance (vigilance decrement), whereas a positive slope denotes increasing engagement. We also calculated the within-trial delta (Δ), the difference in orienting success rate between the first- and last-time bins, where a positive Δ indicates a decline in performance. Both measures were analysed across stripe widths (Figure 3G–H, Figure S3C–D) and, to assess overall within-trial change, after averaging each larva’s values across stripe widths (Figure 3I–J, Figure S3E–F).

Under low contrast, neither the within-trial performance slope (Figure 3G; Mixed-effects model, Genotype: F(1, 770) = 2.785, P = 0.0955) or Δ (Figure 3H; Mixed-effects model, Genotype: F(1, 770) = 2.54, P = 0.1110) differed significantly between genotypes, and there was no effect of stripe width or a genotype × stripe width interaction (all P > 0.24). However, when each larvae’s values when averaged across all stripe widths, mutants were seen to have a more negative within-trial slope than controls (Figure 3I; mutants: −0.0036 ± 0.0011; controls: 0.0005 ± 0.0011; unpaired t-test, t(131) = 2.51, P = 0.013) and a larger positive Δ (Figure 3J; mutants: 0.045 ± 0.014; controls: −0.003 ± 0.014; unpaired t-test, t(131) = 2.42, P = 0.017). Under high contrast, there was no genotype difference in either the slope (Figure S3C; Mixed-effects model, Genotype: F(1, 131) = 0.02, P = 0.886; Figure S3E; Mann-Whitney U test, U = [2150], P = [0.81, confirm]) or Δ (Figure S3D; Mixed-effects model, Genotype: F(1, 131) = 0.06, P = 0.809; Figure S3F; Mann-Whitney U test, U = 2184, P = 0.926), and again no effect of stripe width or interaction (all P > 0.41). Together, these results highlight that under low contrast, *adgrl3.1* mutants show a decline in the tracking performance over the course of each trial. While this was too small to detect at an individual strip width, there was a consistent behaviour across them, which was absent under a high contrast.

In summary, our result show that the closed-loop virtual reality system allows for a high-resolution understanding of *adgrl3.1*-associated hyperlocomotion, establishing that increased locomotor output is driven by shorter, more stereotyped interbout intervals and extended bout lengths, with a decline in tracking performance under low contrast. This approach provides a richer and more precise phenotypic profile of ADHD-candidate risk genes than standard locomotion assays.

## Discussion

In this study, we developed a closed-loop virtual reality paradigm to explore the behavioural consequences of *adgrl3.1* loss in larval zebrafish, providing the first high-resolution phenotyping of this ADHD-linked gene within a stimulus-driven context. Consistent with previous studies reporting increased locomotor activity in *adgrl3.1^-/-^*mutants (Fontana et al., 2023; Regan et al., 2021; Sveinsdottir et al., 2023), CRISPant mutant larvae travelled greater distances than controls during visual stimulation. Furthermore, larval swimming performance followed the expected Gaussian tuning relative to stripe width, peaking at approximately 10.3 mm, in agreement with previous reports (Reynolds et al., 2026). However, by combining controlled visual stimulation with fine-scale locomotor analysis, our closed-loop VR paradigm extends beyond conventional gross hyperactivity measures, revealing that increased output is driven by specific alterations in temporal organization, bout structure, and motor variability.

At a gross level, *adgrl3.1* mutants travelled greater distances than controls, in both low- and high-contrast conditions. Crucially though, this increase does not arise from a uniform increase across all motor parameters. Instead, hyperactivity was driven by a combination of shorter IBIs and longer bout lengths, resulting in higher frequency of swim bouts. In contrast, swim rate did not differ significantly between genotypes, and differences in bout duration were limited. These findings suggest that *adgrl3.1* disruption affects the temporal and spatial organisation of locomotion rather than uniform changes across all measured motor parameters.

IBI was the most sensitive metric, showing consistently reduced duration and variability across visual conditions. While control larvae exhibited flexible IBI patterns, mutants instead produced more frequent and consistent bout timings. This pattern is compatible with a more stereotyped organisation of motor output and may reflect impaired inhibitory control and elevated baseline arousal, aligning with models of impulsivity and dysregulated noradrenergic signalling in ADHD (Biederman & Spencer, 1999; Nigg, 2001; Sigurdardottir et al., 2021). However, reduced variability may also suggest diminished behavioural flexibility to changing environmental demands. Given that IBIs are linked to the integration of sensory input with internal state (Johnson et al., 2020), these findings suggest that *adgrl3.1* disruption alters this integration, promoting high-frequency, less variable responses. Importantly, reduced variability alone does not demonstrate impaired behavioural flexibility and future experiments that quantify behavioural transitions under changing stimulus conditions will be important for determining whether the observed consistency reflects reduced flexibility or another alteration in motor regulation.

This altered temporal regulation was also evident during the pre-stimulus black screen phase. Baseline analysis revealed that *adgrl3.1* mutants exhibited significantly shorter IBIs prior to visual motion exposure, while overall speed, bout distance, and event frequency remained comparable. This selective baseline difference points to an intrinsic increase in motor excitability and a lower threshold for initiating motor events, independent of immediate visual feedback.

Mutant larvae also displayed longer swim bouts, with the largest difference observed under low-contrast conditions. This suggests that the loss of *adgrl3.1* alters the regulation of visuomotor behaviour, where impaired visual discrimination may contribute to this motor output. In ADHD, there notably can be dysregulated effort allocation, producing either over- or under-engagement depending on task demands. In addition, mutants exhibited lower behavioural variability through reduced coefficient of variation in both IBI and average swimming speeds than controls. Together with the shorter IBIs observed, this reduced variability suggests a more consistent temporal organisation. Altered trial-to-trial variability has been noted in some human ADHD studies, particularly in motor tasks, and may be a signature of dysregulated internal state modulation (van der Meer et al., 2016). By combining controlled visual stimulation with fine-scale locomotor analysis, the closed-loop VR paradigm therefore extends beyond conventional understanding of hyperactivity, enabling the investigation of how genetic mutations influence the organisation of motor behaviour across different sensory conditions.

Together, these findings extend the characterisation of *adgrl3.1* mutants beyond gross hyperactivity to reveal distinct fine-scale behavioural changes of shorter interbout intervals, longer bout lengths, and reduced variability, indicating that increased activity arises from specific changes in how locomotor activity is organised. These alterations suggest that increased locomotor output is accompanied by changes in the temporal structure and consistency of swimming, rather than representing a uniform increase in activity across all behavioural parameters. These phenotypes may be suggestive of ADHD-associated processes such as impulsivity, reduced flexibility, and altered sensory–motor coupling, although these relationships remain to be confirmed. Nevertheless, altered movement timing and reduced variability provide more detailed quantifiable phenotypes for investigating how genetic perturbations influence motor regulation and sensorimotor behaviour than standard locomotor assays. By resolving hyperactivity into distinct behavioural components under controlled visual stimulation, the closed-loop VR paradigm offers a more detailed framework for investigating the behavioural consequences of ADHD-associated genetic variation, and for distinguishing phenotypes that may appear similar when assessed using conventional locomotor measures alone.

Importantly, interpretation of these findings would benefit from further validation of the genetic model. As the experiments were performed with F0 CRISPants, mosaic editing and variation in the extent of gene disruption may contribute to phenotypic variability. Although genotyping supports successful targeting of the *adgrl3.1 locus*, further validation through mRNA or protein detection would confirm the loss of *adgrl3.1*. More broadly, establishing how these larval phenotypes relate to the complex cognitive and behavioural features of ADHD will require further investigation. The present assay provides a controlled way to quantify how locomotor behaviour is organised under behaviour-contingent visual feedback. Therefore, future work combining this assay with pharmacological perturbation, circuit imaging, or explicitly validated behavioural tasks could help understand how the neural mechanisms underlying the behavioural changes map onto specific ADHD-relevant processes.

In conclusion, we extend the characterisation of *adgrl3.1*-associated hyperlocomotion to show that it emerges from shorter and more stereotyped IBIs, combined with longer bouts, and reduced variability. These findings highlight the importance of segmenting locomotor output into its constituent parameters rather than treating hyperactivity as a single behavioural measure. These alterations in behaviour may mirror core ADHD phenotypes of impulsivity and reduced behavioural flexibility. By combining genetic manipulation with fine-scale behavioural quantification, this approach addresses critical gaps in our understanding of ADGRL3 function and also establishes a scalable platform for drug discovery and mechanistic dissection of neurodevelopmental disorders.

## Acknowledgements

This work was supported by the MRC (MR/W006251/1, PhD Studentship to ER; MRC DTP MR/N013700/1 to PR) and The NC3R’s (NC/X001520/1 to RH and CD-E). We thank members of the team for their insightful discussions.

## Author Contributions

RH and PR conceived the overall study and designed the experiment. ER created the F0 CRISPant mutant animals, PR performed all experiments and analysed the data with input from CD-E. PR wrote the first draft of the manuscript with input from all authors.

## Data availability

Raw data can be found on the Hindges Lab Github Page (github.com/Hindges-Lab).

## Code availability

The computer code used to analyse data is available from the Hindges Lab Github page (github.com/Hindges-Lab).

