## Supplemental Figures and Table for "Virtual reality reveals fine-scale alterations in behaviour following loss of the ADHD-linked gene adgrl3.1 in zebrafish"

A)

B)

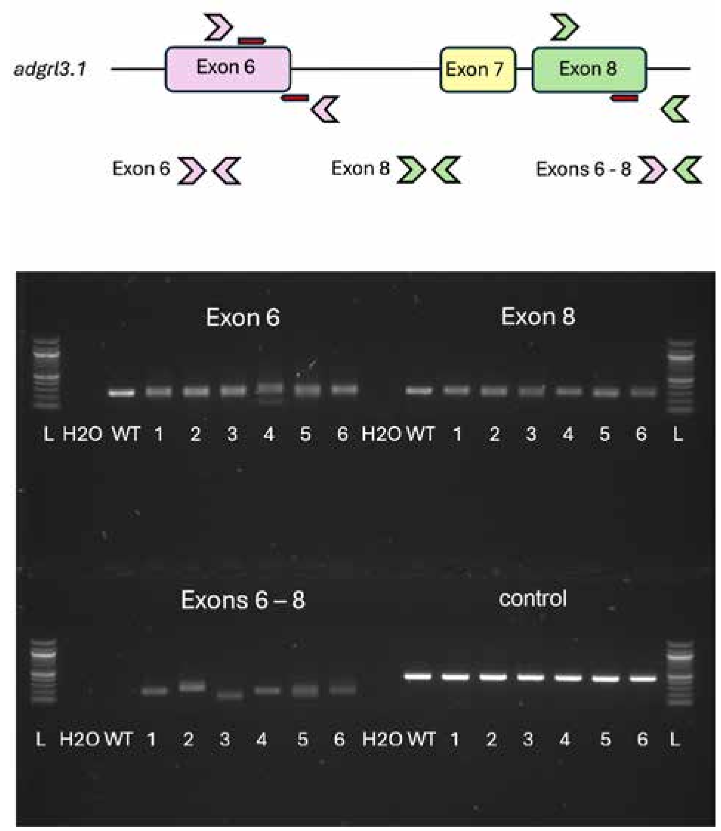


**Supplemental Figure S1. PCR-based characterisation of *adgrl3.1* CRISPants.**

**A)** PCR approach with location and direction of gRNAs (red arrows), and location and direction of primers (large pink and green arrows), on each exon of *adgrl3.1*. **B)** Example of genotyping results showing indel mutations for all individual larvae in exon 6, likely indels in exon 8, and partial excisions between exons 6–8. For each sample a positive control was done using a primer pair in an unrelated genomic location (tenm4 gene). H2O, negative water control; L, DNA ladder; WT, DNA from uninjected wild-type animals.

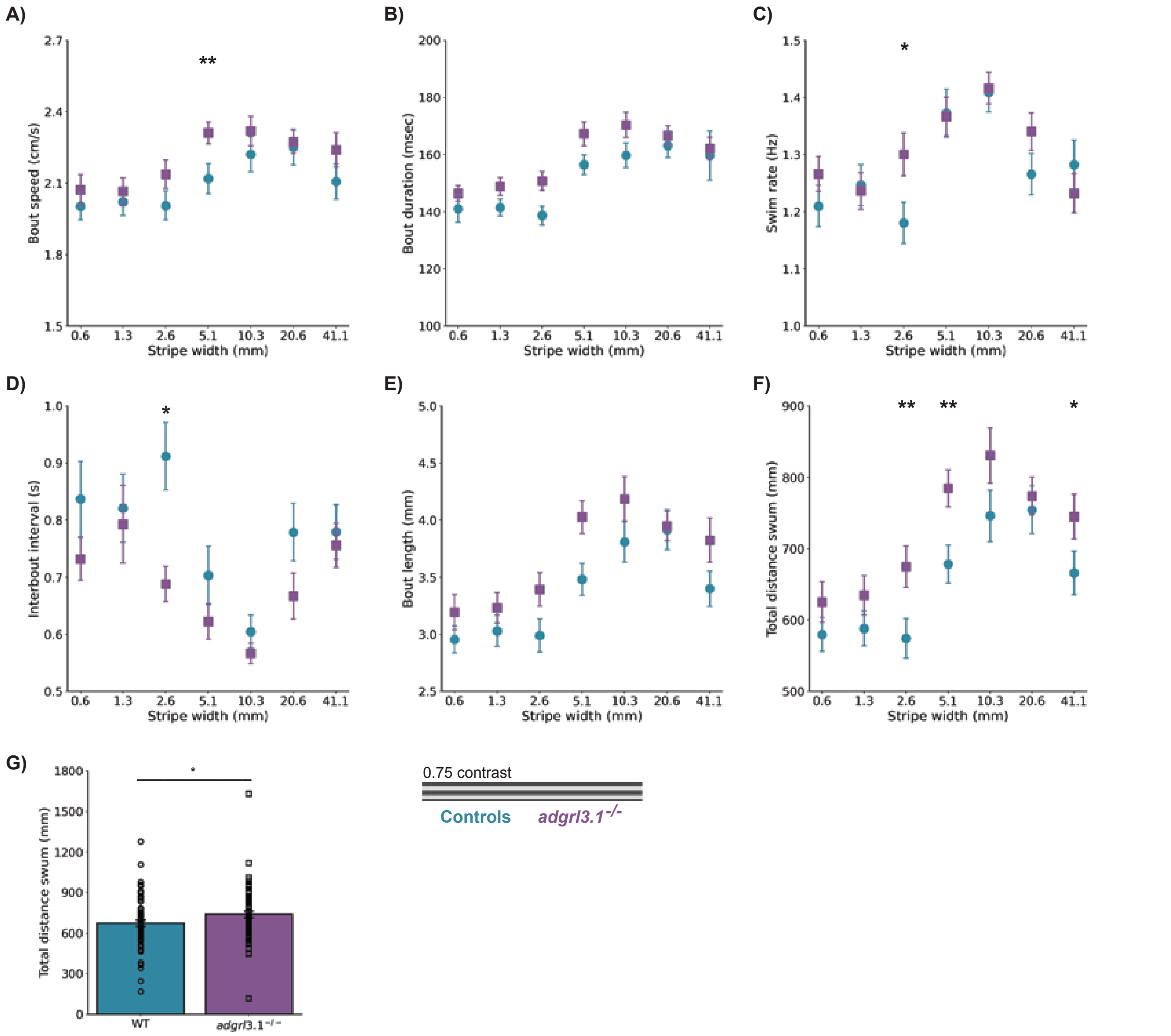


**Supplemental Figure S2. Increased locomotor output in *adgrl3.1* mutants viewing a higher contrast stimulus is driven by altered bout structure.**

**A-F)** Various swim behaviours measured across stripe widths at a 0.75 AU contrast condition for control larvae (blue), and loss-of-function *adgrl3.1* mutants (purple). **A)** Average bout speed (Mixed-effects model, Genotype: P = 0.1675; Stripe width: ***P < 0.001; Stripe width at 5.1 mm: **P = 0.0032). **B)** Average bout duration (γ GLMM, Genotype: P = 0.207; Stripe width: ***P < 0.001. **C)** Swim rate measured in bouts per second (Mixed-effects model, Genotype: P = 0.4592; Stripe width: ***P < 0.001). **D)** Interbout interval (IBI) (γ GLMM, Genotype: P = 0.082; Stripe width: ***P < 0.001; Stripe width at 2.6 mm: *P = 0.013. **E)** Bout length (distance travelled per bout) γ GLMM, Genotype: P = 0.169; Stripe width: ***P < 0.001). **F)** Total distance travelled (Mixed-effects model, Genotype: *P = 0.0159; Stripe width: ***P < 0.001; Stripe width at 2.6 mm: **P = 0.0044; Stripe width at 5.1 mm: **P = 0.0039; Stripe width at 41.1mm: *P = 0.0441). **G)** Total distance travelled at a 0.75 AU contrast condition for control larvae (blue), and loss-of-function *adgrl3.1* mutants (purple) (Mann-Whitney U Test, *P = 0.0471). **A–G)** n = 62 for *adgrl3.1* mutant larvae, n = 70 for control larvae. All error bars are mean ± s.e.m. over fish. Asterisks indicate significance *P < 0.05, **P < 0.01.

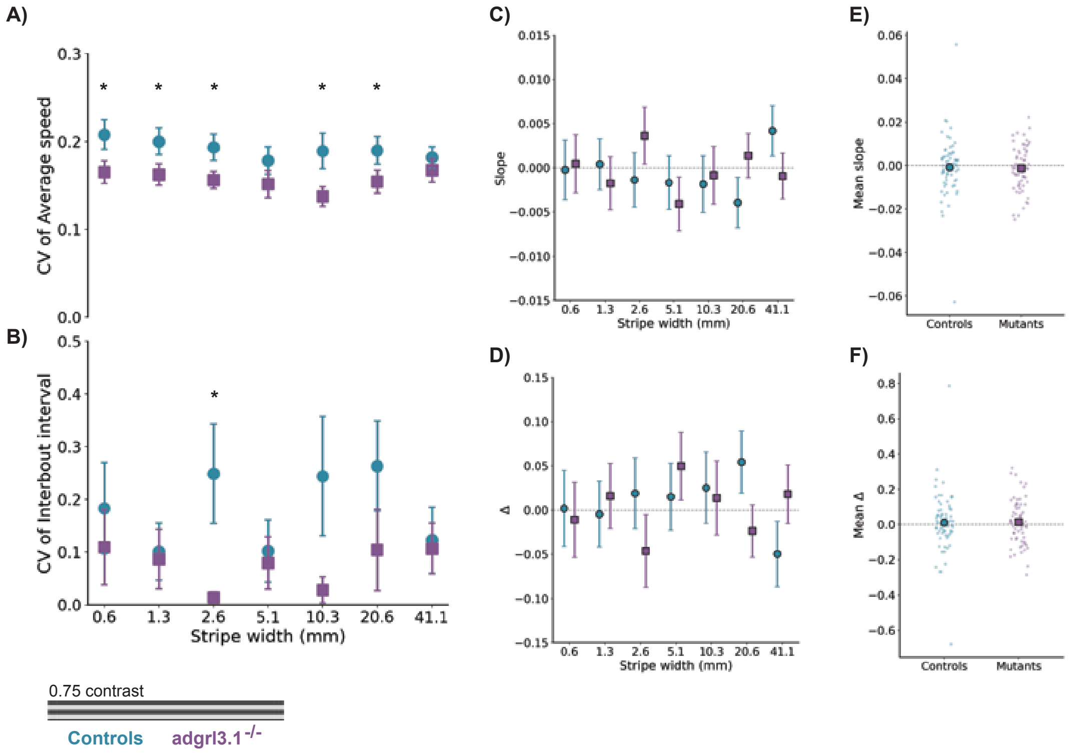


**Supplemental Figure S3. Reduced behavioural variability altered baseline interbout interval are present in *adgrl3.1* mutants.**

**A)** Coefficient of variation (CV) for average speed across stripe widths to indicate flexibility in motor output (Mixed-effects model, Stripe width: P = 0.2736; Genotype: **P = 0.0067; Stripe width 0.6 mm: *P = 0.0288; Stripe width 1.3 mm: *P = 0.0158; Stripe width 2.6 mm: *P = 0.0433; Stripe width 10.3 mm: *P = 0.0229; Stripe width 20.6 mm: *P = 0.0235). **B)** Coefficient of variation (CV) for interbout interval (IBI) across stripe widths to indicate flexibility in motor output (Mixed-effects model, Stripe width: P = 0.8476; Genotype: P = 0.0533; Stripe width 2.6 mm: *P = 0.0161). **C)** Within-trial slope of successful optomotor responses over stripe widths. Negative slopes indicate declining performance (vigilance decrement). (Mixed-effects model, Genotype: P = 0.8859; Stripe width: P = 0.7671). **D)** Within-trial delta (Δ) in response success between early and late trial phases across stripe widths, where larger positive deltas indicate decline in performance over trials (Mixed-effects model, Genotype: P = 0.8094; Stripe width: P = 0.8423). **E)** Within-trial slope averaged across stripe widths for each larva (Mann-Whitney U test, U = 2129, P = 0.8513). **F)** Within-trial Δ averaged across stripe widths for each larva (Mann-Whitney U test, U = 2184, P = 0.9258). **A-F)** n = 62 for *adgrl3.1* mutant larvae, n = 70 for control larvae. All error bars are mean ± s.e.m. over fish. Asterisks indicate significance *P < 0.05.

**Supplemental Table 1: Sequences for PCR primers and gRNAs for *adgrl3.1* CRISPants**

| | |
|---|---|
| Exon 6 forward primer | CTCCCGTCACTGTTGACTATTG |
| Exon 6 reverse primer | CAGAGAGCAATGCACATACACA |
| Exon 8 forward primer | GCTGAACAGACAAAGGGAGAGT |
| Exon 8 reverse primer | TGAGCTATCTCATGCAAAAGGA |
| Exon 6 gRNA 1 | CTGCTGAAATCTCCTGGCCA CGG |
| Exon 6 gRNA 2 | TACTAACCTATGGTGCCAGG TGG |
| Exon 8 gRNA | GGCGGTGTCTTTTCCCCCAG GGG |
